# Machine learning enables high-throughput, low-replicate screening for novel anti-seizure targets and compounds using combined movement and calcium fluorescence in larval zebrafish

**DOI:** 10.1101/2024.08.01.606228

**Authors:** Christopher Michael McGraw, Annapurna Poduri

## Abstract

Identifying new, more efficacious anti-seizure medications (ASMs) is challenging, partly due to limitations in animal-based assays. Zebrafish (*Danio rerio*) can serve as a model of chemical and genetic seizures, but methods for detecting seizure-like activity in zebrafish, though powerful, have been hampered by low sensitivity (locomotor/behavioral assays) or low-throughput (tectal electrophysiology or calcium fluorescence microscopy). To address these issues, we developed a novel approach to assay seizure-like activity using combined locomotor and calcium fluorescence features, measured simultaneously from unrestrained larval zebrafish using a 96-well fluorescent plate reader. Using custom software to track fish movement and changes in fluorescence (deltaF/F0) from high-speed time-series (12.6Hz), we trained logistic classifiers using elastic net regression to distinguish seizure-like activity from non-seizure related changes based on event-specific and subject-specific features in response to the GABA_A_R antagonist, pentylenetetrazole (PTZ). We demonstrate that a classifier trained on combined movement and fluorescence data achieves high accuracy (“PTZ M+F”; area-under-curve receiver-operator characteristic (AUC-ROC): 0.98; F1 score: 0.912) and out-performs classifiers trained on movement (“PTZ M”; AUC-ROC: 0.9, F1: 0.9) or fluorescence features alone (“PTZ F”; AUC-ROC 0.96; F1: 0.87). The rate of classified seizure-like events increases as a dose-response to PTZ (serial dose escalation, 0, 2.5mM, 15mM) and is strongly suppressed by ASM treatment (valproic acid, VPA; tiagabine, TGB). At high-dose PTZ, we show that VPA reduces seizure-like activity defined by either “PTZ M+F” or “PTZ M” classifiers. Meanwhile, TGB selectively reduces events defined by the “PTZ M+F” classifier, paralleling previous reports that TGB reduces electrographic but not locomotor seizures and highlighting the potential for our approach to combine features of previously orthogonal assays. Using ASM benchmark data, we employ bootstrap simulation to calculate the expected statistical power of our method as a function of sample size. We demonstrate that anti-seizure responses (robust strictly standardized mean difference, RSSMD, versus control) with magnitudes similar to those associated with VPA or TGB can be reliably detected (true positive rate (TPR) > 90%) with as few as N=4 biological replicates per group, while maintaining a 5% false positive rate. In a prospective test screen with 3-6 replicates per group and on-plate controls, the anti-seizure effect of 4 out of 5 tested ASMs (CBZ, LEV, LZP, TGB) was detected. In summary, we demonstrate a simple high-throughput approach to whole organism anti-seizure phenotyping combining two previously reported metrics to facilitate screens for novel anti-seizure interventions in zebrafish.

## 1. Introduction

Epilepsy is a chronic condition comprised of spontaneous recurrent seizures that affects more than 70 million people worldwide^1^, roughly 30% of whom remain poorly controlled after two therapeutic trials of anti-seizure medications (ASMs)^1^. The search for novel ASMs continues, particularly for drug-resistant epilepsy (DRE)^2^, as the introduction of novel ASMs in the last 30 years has not substantially improved rates of DRE^1^. Although some recent efforts to rationally design novel target-based therapies^3^, drug screening using functional assays of seizure-like activity remain both the backbone and a major bottleneck of ASM discovery^2^. Traditionally based on classic rodent models of acute seizures^1^, current efforts have expanded to employ newer animal models such as zebrafish^4,5^, in addition to genetically modified animals (rodents and zebrafish)^1,2^ and cellular models of epilepsy, including human neuronal cultures from patient-derived induced pluripotent stem cells (iPSCs)^7^ .

Zebrafish in particular has shown great promise as a model of chemically induced seizures (for example in response to PTZ^4^) including drug-refractory seizures (allyl-glycine^8^), and as a model of seizures related to genetic epilepsy (*scn1lab*^9^; *stxbp1b*^10^; *pcdh19*^11^, and many others^12,13^), with excellent predictive validity. In fact, the novel ASM fenfluramine^14^ was identified from a screen in the *scn1lab* zebrafish (a model of Dravet syndrome) and has shown efficacy in reducing seizures in individuals with Dravet syndrome across two randomized, placebo-controlled clinical trials^15,16^.

Assays for seizure-like activity in larval zebrafish are usually conducted from 3-7 days post fertilization (d.p.f.) and commonly include: locomotor assessment^4,14^ to detect seizure-related bouts of rapid swimming and increased distance travelled; electrophysiology^5,12,13^ to identify epileptiform discharges in local field potentials recorded from optic tectum; and calcium fluorescence to detect high amplitude synchronized neuronal depolarization usually in the optic tectum^17,18^ or whole brain^19,20^ using confocal or single plane illumination (“lightsheet”) microscopy. In cases where the ASM responses in two assays have been studied in parallel, high concordance has been reported^5^, although it has been noted that locomotor assays may be less sensitive to detect the response of some antiseizure drugs (for example tiagabine, TGB^5^) or to detect seizure-like activity in genetic models^12^. Meanwhile, although assays based on electrophysiology and calcium fluorescence microscopy appear more sensitive, they are considerably lower throughput. Last, quantification of seizure-like activity associated with proconvulsants that also cause immobility (such as organophosphate toxins, like DFP^21^) or seizure-like activity not associated with rapid swimming (as reported in several genetic models^12,13^) may be wholly inaccessible by locomotor assays, requiring the use of electrophysiology or microscopy. A method that could combine the sensitivity of neurophysiological approaches such as tectal EEG or calcium fluorescence microscopy with the throughput of locomotor-based assays would therefore significantly enhance existing processes for the discovery of novel antiseizure interventions in larval zebrafish.

Here we report our efforts to develop a higher-throughput approach using a fluorescent plate reader and machine learning to assess combined movement- and fluorescence-based measures of seizure-like activity from larval zebrafish expressing neuronal GCaMP6s. We show that this method detects seizure-like activity induced by the GABA_A_ receptor antagonist pentylenetetrazole (PTZ) with high accuracy, and can quantify the response to ASMs. We show how combined movement and fluorescence profiling distinguishes differential effects of TGB, thus reconciling conflicting data in the literature derived from dedicated locomotor versus electrographic assays. We benchmark our assay using known ASMs (VPA, TGB) and use bootstrap simulations to establish screening thresholds based on the robust strictly standardized mean difference (RSSMD^22^) in rate of PTZ-related seizure-like activity between treated versus control. These simulations support screens based on relatively low number of biological replicates (N=4-8), and we demonstrate the ability to detect the anti-seizure effect of several known ASMs using these thresholds.

Together, our method is a simple and powerful higher throughput approach to whole organism phenotyping of seizure-like activity that should facilitate future screens to identify novel anti-seizure compounds and targets.

## 2. Materials and methods

### Zebrafish maintenance

GCaMP6s^23^ zebrafish (*Danio rerio*) with *nacre* pigmentation phenotype used for all experiments were obtained as TG(*elav3*::Gcamp6s); *mitfa*^w2/w2^ (abbreviated, *GCaMP6s*; generous gift from Florian Engert, Harvard University). Zebrafish were maintained by in-crosses, and larvae periodically selected for “high” GCaMP6s expression based on epifluorescence. All fish were maintained on a 14H:10H day-night cycle. All procedures were approved by BCH Animal Welfare Assurance (IACUC protocol #00001775).

### Compound preparation

Test compounds included the following: valproic acid (Sigma, P4543), tiagabine (TCI, T3165), carbamazepine (Sigma, C4024), phenytoin (5,5-diphenylhydantoin; Millipore Sigma; D4007), lorazepam (Sigma, L1764), ampicillin (Sigma, A9393), and caffeine (Sigma, C0750). Test compounds were prepared in DMSO (0.1% final concentration), except VPA which was prepared in fish water.

### PTZ dose escalation

PTZ (Sigma; stored -20degC) was prepared fresh in sterile fish water (Instant Ocean) to a stock concentration 26mM, then diluted to intermediate concentrations. For serial dose escalation experiments, a standard 10uL volume from PTZ Stock 1 (25.7mM) was added to each well of a 96-well plate (100uL starting volume per well) to yield 2.5mM, followed by an additional 10uL from PTZ Stock 2 (152.5mM; 110uL starting volume per well) to yield 15mM final concentration (final well volume, 120uL). For experiments involving anti-seizure drug pretreatment, anti-seizure drugs were administered in a standard 10uL volume. Following baseline recording, a standard 10uL volume from PTZ Stock 1 (30mM) was added to each well of a 96-well plate (110uL starting volume per well) to yield 2.5mM, followed by an additional 10uL from PTZ Stock 2 (165mM; 120uL starting volume per well) to yield 15mM final concentration (final well volume, 130uL). Pipetting was performed manually with a multi-channel pipettor. Three sequential 30-minute recordings were performed during baseline, PTZ 2.5mM, and PTZ 15mM conditions, respectively.

### Calcium fluorescence imaging

Individual unrestrained larval zebrafish (dpf 5) are placed into wells of an optical 96-well plate (Greiner 655076) in 100uL sterile fish water (Instant Ocean) and imaged using the FDSS7000EX fluorescent plate reader (Hamamatsu; software version 2). Specimens are illuminated by a Xenon light source passed through a 480nm filter. Epifluorescence from below the specimen is filtered (540nm) and collected by EM-CCD, allowing all wells to be recorded simultaneously. Data is collected as 256x256, 16-bit image at ∼12.6 Hz (79 msec interval), 2x2 binning, sensitivity setting = 1. Image data was extracted from the .FLI file using ImageJ or MATLAB based on the following parameters: 16-bit unsigned, 256x256, offset 66809, gap 32 bytes. Raw fluorescence videos from baseline (**Supplementary Video 1**) and during PTZ 15mM trial (**Supplementary Video 2**) are provided. Analysis was performed in MATLAB to extract position, linear and angular velocity, and changes in calcium fluorescence using a moving average deltaF/F0 method.

### Analysis of calcium fluorescence data

An algorithm to track changes in calcium activity using a “moving delta F/F0” was devised in MATLAB. The initial 256x256 time-series is segmented into individual wells (∼14 x 14 pixels, ∼0.513mm per pixel) based on a pre-specified plate map. For each well, the *n* x *m* x *t* time-series is expanded to 2*n* x 2*m* x *t* using bicubic interpolation before further processing. Calcium transients are detected based on the normalized instantaneous average fluorescence for the area of the fish body within the well by the following formula: (average F_fish_(t) - F_0_)/F_0_, where F_0_ = average Ffish (averaged over each pixel, for each time sample), and smoothed with a 1000-sample (∼79 seconds) boxcar moving average. Fish x,y position is tracked based on a weighted centroid, and linear and angular velocity estimated. The minimum detectable change in the position of a larval zebrafish is estimated to be 0.256mm, corresponding to the size of one pixel after interpolation. For detecting significant fluctuations in calcium fluorescence (referred to as calcium events), the F/F0 time-series is further smoothed with a 25-sample (∼1.975 seconds) boxcar moving average. Calcium events are initially detected from the smoothed delta F/F0 time-series by identifying peaks that exceed an empirically determined permissive threshold (0.05), while the start and end of each event is identified by the zero-crossing of the smoothed 1st derivative.

Subsequently, multiple per-event measurements are obtained for each event based on combined movement and fluorescence measurements, including: (1) MaxIntensity_F_centroid: the maximum fluorescence value of the detected fish during an event; (2) MaxIntensity_F_F0_centroid: the maximum delta F/F0 value within the boundary of the detected fish during an event; (3) distance_xy_mm: total distance moved during an event in millimeters; (4) duration_sec: elapsed time in seconds; and (5) total_revolutions: number of complete circles traveled by the fish during an event.

In addition, multiple per-fish measurements are obtained, including: (1) maxRange_2: the maximum fluorescence value observed during the recording; (2) totalCentroidSize_mode_mm2: the total area in square millimeters of the detected fish that exceeded a hard-coded threshold above sensor noise, which relates to the brightness of the fish.

### UMAP visualization

A single manifold approximation and projection was performed in R using function *umap* (R package, *umap*) and the combined datasets of calcium events from experiments reported here and another experiment involving genetically modified zebrafish. Briefly, the dataset comprised of 67366 unique calcium events from 395 unique animals (rows = calcium events, columns = variables) was filtered to include only the following columns: MaxIntensity_F_F0_centroid; distance_xy_mm; maxVeloc_xy_mm_sec; duration_sec; timeSpentMoving_xy_sec_prct; totalRevolutions_perEvent; and totalCentroidSize_mode_mm2_round. Each column was centered and scaled prior to calculating UMAP. Visualization was performed with function *ggplot* (R package, *ggplot2*); a 2D kernel density estimation was applied using geom_density_2d(contour_var = “ndensity”), which scales the density estimate to a maximum of 1 within each facet panel. To facilitate comparison, all visualizations shown employed the same embedding.

### Supervised machine learning for event classification

To differentiate calcium events related to seizure-like activity from other causes of calcium fluctuation, a logistic classifier was fit to a combination of per-event measurements and per-fish measurements using elastic net regression using glmnet (R package, *glmnet*) and train (R package, *caret*) using R.

Training data were obtained from serial PTZ dose-escalation trials (N=64 fish, >4000 events). Fish lacking minimum fluorescence criteria (mode of fluorescence area < 0.05 mm^2^) were excluded from analysis. Trial labels were used in lieu of manual labeling of individual events. All detected events during the baseline trial period (PTZ=0; 30 minutes) were used for “non-seizure activity” while all events during the subsequent PTZ trial period (PTZ=15mM; 30 minutes) was used for “seizure-like activity”. Data were divided into 70:30 train:test split, with 10-fold cross validation, alpha range: 0,0.5, 1, lambda range: 0.1, 1, 10, and metric = “accuracy”. Model performance was evaluated using package MLeval.

Three models were fit as below: 1) Movement and fluorescence (“M+F”): Event data was pre-filtered for high fluorescence (MaxIntensity_F_F0_centroid >0.1). An interaction term was used to account for possible interactions between event-based measurements and fish-based measurements. The model formula: Conditions_names ∼ (MaxIntensity_F_centroid + MaxIntensity_F_F0_centroid + distance_xy_mm + duration_sec) * (maxRange_2 + totalCentroidSize_mode_mm2). The chosen model employed alpha = 0, and lambda 0.1. 2) *Movement only*: Event data was used without pre-filtering in order to minimize additional selection based on magnitude of calcium fluorescence, though some selection is unavoidable based on the initial selection in Matlab. The model formula: Conditions_names ∼ (maxVeloc_xy_mm_sec + distance_xy_mm). The chosen model employed alpha =0.5, and lambda 0.1. 3) *Fluorescence only*: Event data was pre-filtered for high fluorescence MaxIntensity_F_F0_centroid >0.1. The model formula was: Conditions_names ∼ (MaxIntensity_F_centroid + MaxIntensity_F_F0_centroid + duration_sec + maxRange_2 + totalCentroidSize_mode_mm2). The chosen model employed alpha = 0, and lambda 0.1.

With the exception of the preliminary PTZ dataset without antiseizure treatment (e.g. Figure 3, and control data in Figure 4), no additional experimental data were present in the training dataset.

### Bootstrap simulations

Bootstrap simulations were conducted in R using the *rep_sample_n()* function (R package, *moderndive*) essentially as described in the main text.

### Statistical analysis

Statistical analysis was conducted in R. Unless otherwise stated, pairwise differences in measured parameters between groups were assessed using the non-parametric unpaired two-sided Wilcoxon rank sum test (aka Mann-Whitney test; R packages, *ggpubr*; function, *compare_means*) with false discovery rate (FDR) correction (Benjamini-Hochberg method) to maintain alpha < 0.05 (R package, *stats*). Only adjusted p-values are reported.

## 3. Results

### 3.1. A fluorescent plate reader is suitable for recording combined calcium fluorescence and locomotor activity from freely swimming larval zebrafish at baseline and in response to pro-convulsant drugs

To develop a high-throughput approach, we first asked whether a conventional fluorescent plate reader would provide sufficient temporal and spatial resolution to detect changes in neuronal calcium fluorescence in response to a proconvulsant (**Fig 1A**). Preliminary testing demonstrated that detection of seizure-like activity in response to PTZ (15mM) was feasible (**Fig 1B**), resulting in large amplitude fluctuations in the measured fluorescence intensity, despite epifluorescence measurements being collected from the ventral aspect of unrestrained larval fish.

**Figure 1.**
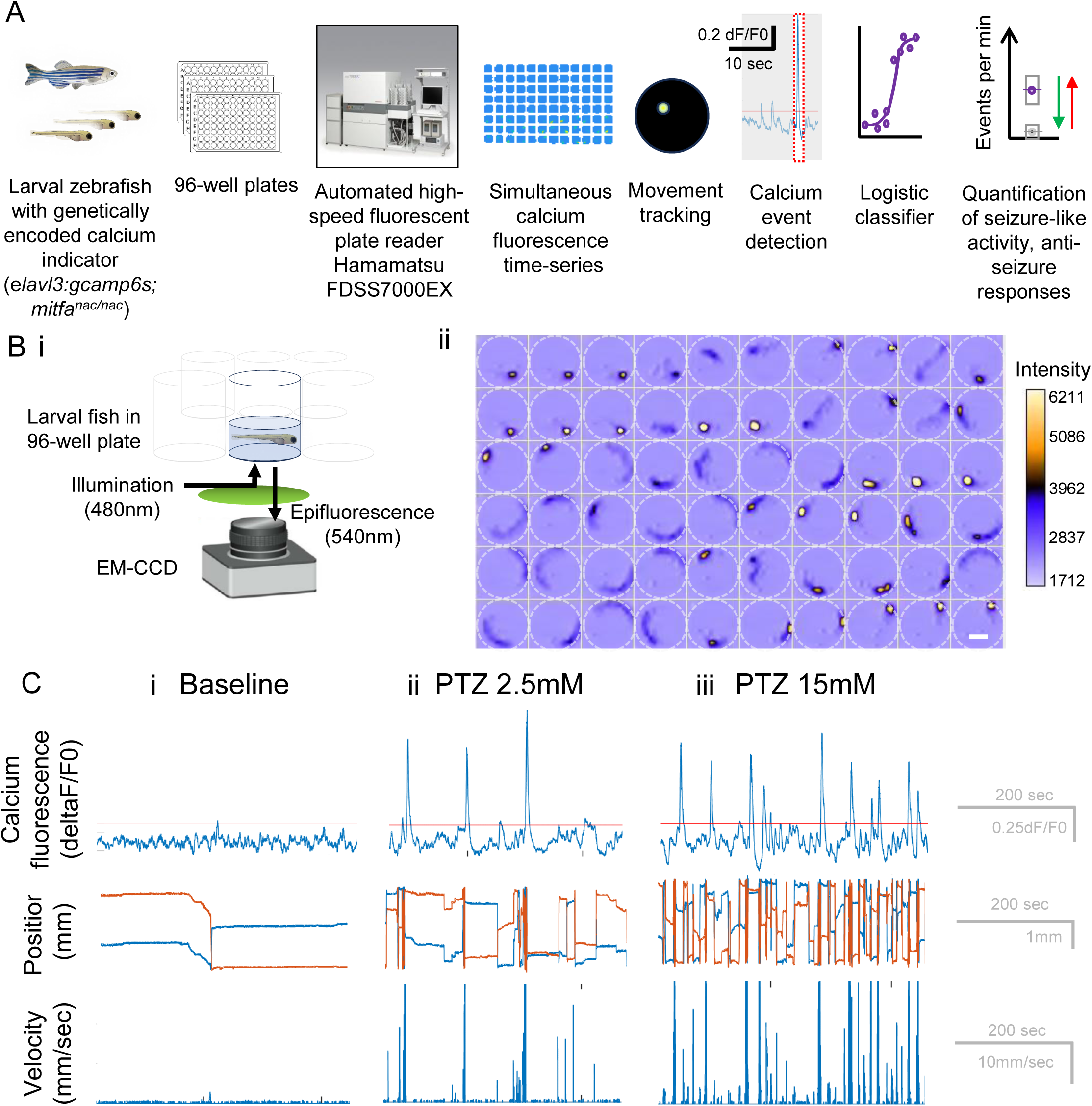
The FDSS7000EX is suitable for recording combined movement and calcium fluorescence time-series from unrestrained GCaMP6s larval zebrafish at baseline and in response to pro-convulsant drugs. (A) Schematic for anti-seizure profiling using calcium fluorescence from larval zebrafish in 96-well format on the FDSS7000EX fluorescent plate reader. (B) Schematic of light-path for epifluorescence recording (*i*) and representative time-series montage (*ii*) of a single larvae over time after incubation with proconvulsant (15mM PTZ). Time = 79msec between frames. Dotted circle shows position of the well. Scale bar in last frame is 3mm. (C) Combined estimation of change in calcium fluorescence (deltaF/F0; *upper*), fish position (*middle*), and velocity (*lower*) at baseline (*left*), low-dose PTZ (2.5mM, *center*), and high-dose PTZ (15mM, *right*). No event classification has been applied. Red reference line marks the position of dF/F0 = 0.1.

We next performed a PTZ serial dose escalation trial (0, 2.5, 15mM) and demonstrate that measured fluorescence throughout the recording has sufficient spatial resolution to track position and velocity in the 2D plane, and sufficient temporal resolution (∼12.6Hz) to show characteristic changes in normalized fluorescence (dF/F0) that resemble seizure-like activity in these unrestrained larvae (**Fig 1C**). From representative examples, during baseline conditions (PTZ=0), we note some rare changes in calcium fluorescence related to movement (**Fig 1Ci**). With PTZ, we observe progressive changes in fish activity, with an increasing number of movement episodes with higher velocity (up to 30mm/sec) with increasing PTZ dose. Calcium events (threshold dF/F0 > 0.05) increase in rate and magnitude of delta F/F0 (up to 1 arbitrary unit (a.u.)) during PTZ=2.5mM (**Fig 1Cii**). – values that appear rare during baseline testing. During PTZ=15mM (**Fig 1Ciii**)., the rate of calcium events increases further though the magnitude of delta F/F0 appears to reduce slightly (0.2-0.8 a.u.). These patterns are strongly reminiscent of similar observations regarding dose-dependent increases in the rate of dF/F0 fluctuations derived from whole-brain imaging of PTZ-induced seizures using confocal microscopy in optic tectum of larval zebrafish^17^, suggesting that our simple approach may nevertheless yield similar estimates of seizure-like activity.

We next examined the distribution of data across all calcium events for each group in the PTZ dose-escalation trial with respect to fluorescence- and movement-related measures using the unified manifold approximation and projection (UMAP)^24^ dimensionality reduction approach (**Fig 2A-F**) followed by group-level quantification of event measures after averaging within each larvae (**Fig 2G-K**).

**Figure 2.**
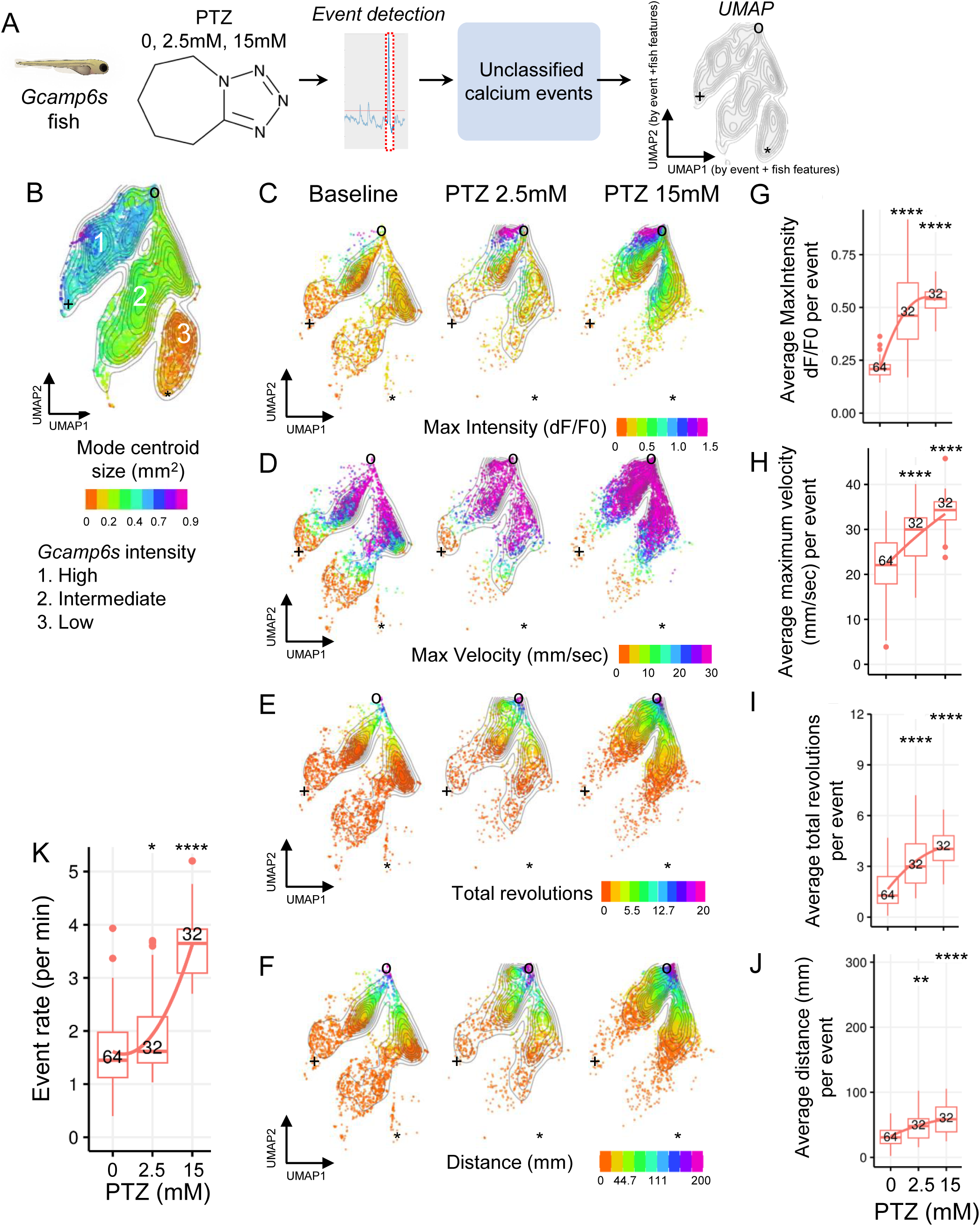
Major features of calcium events from combined movement and fluorescence profiling during PTZ exposure. (**A**) Overview of PTZ dose escalation paradigm (**B**) Event clusters in UMAP space are driven strongly by fish brightness (“mode centroid size”). Clusters 1-3 represent “high Gcamp6”, “intermediate GCaMP6s”, and “low GCaMP6s”. UMAP events are from PTZ-VPA, PTZ-TGB, scn1lab data sets (375 larvae; 67366 calcium events). (**C-J**) UMAP clustering of individual calcium events (C-F) and quantification at the group-level (G-J) following averaging at the fish level. UMAP data are colored according to corresponding color scales for calcium fluorescence intensity (C), maximum event velocity (D), total revolutions (E) and distance travelled per event (F). Fiducial markers (+,o,*) relate to the common event embedding for all data reported in this study. Boxplots are Tukey format, the number of larvae per group is indicated as numerals. (**K**) Rate of calcium events increases in dose-response to PTZ. Statistical testing is FDR-adjusted pair-wise Wilcoxon rank sum test. *, adj P<0.05. **, adj P<0.01. ***, adj P<0.001. ****, adj P<0.0001.

We confirm that GCaMP6s fish show dose-dependent elevations in the max intensity of calcium discharges (dF/F0; **Fig 2C,G**), maximum velocity (**Fig 2D,H**), total revolutions **(Fig 2E,I**), and distance travelled **(Fig 2F, J)**. In addition, the rate of calcium events increases in a step-wise manner (**Fig 2K**).

These analyses revealed several challenges to a naïve approach to develop an assay of seizure-like activity using combined movement and fluorescence measures. First, although PTZ-related events show clear dose-dependent elevations at each level of PTZ across multiple measures, the raw rate of calcium events does not reflect any special knowledge about the features of PTZ-related activity, as reflected by the high baseline rate of calcium events, only slightly lower than those measured at low-dose PTZ. Second, we find that despite efforts to maintain high GCaMP6s brightness among progeny and removing fish below threshold, GCaMP6s fish still display at least 3 levels of brightness (**Fig 2B**), as demonstrated by UMAP plots of fish events colored by fish centroid size above sensor threshold (similar to direct measures of fish fluorescence, **Supplementary Fig 2)**. Fish brightness may impact the signal-to-noise ratio (SNR) for detecting changes in calcium fluorescence (although we could not confirm systematic differences in the SNR of normalized dF/F0 or maximum dF/F0 as a function of fish brightness; **Supplementary Fig 3**), which could make the ability to define properties of seizure-like activity across heterogeneous groups more difficult.

### 3.2. Elastic net logistic regression classifies events as seizure-like with high accuracy

To address the previously noted challenges in a data-driven manner, we turned to supervised machine learning in the form of elastic net logistic regression^25^ to fit classifiers using multiple per-event and per-fish features with the goal of identifying seizure-like events with high confidence (**Fig 3A**). Logistic regression fits a set of coefficients to a sigmoid response function using a defined set of predictors in conjunction with labelled training data from each of two classes of interest. When applied to novel data, the tuned model or classifier yields predictions from 0-1 which reflect the probability of class membership. Elastic net regression^25^ is an approach to fitting logistic models using penalized maximum likelihood which combines features of ridge regression^26^ and lasso regression^27^ to perform regularization and variable selection.

**Figure 3.**
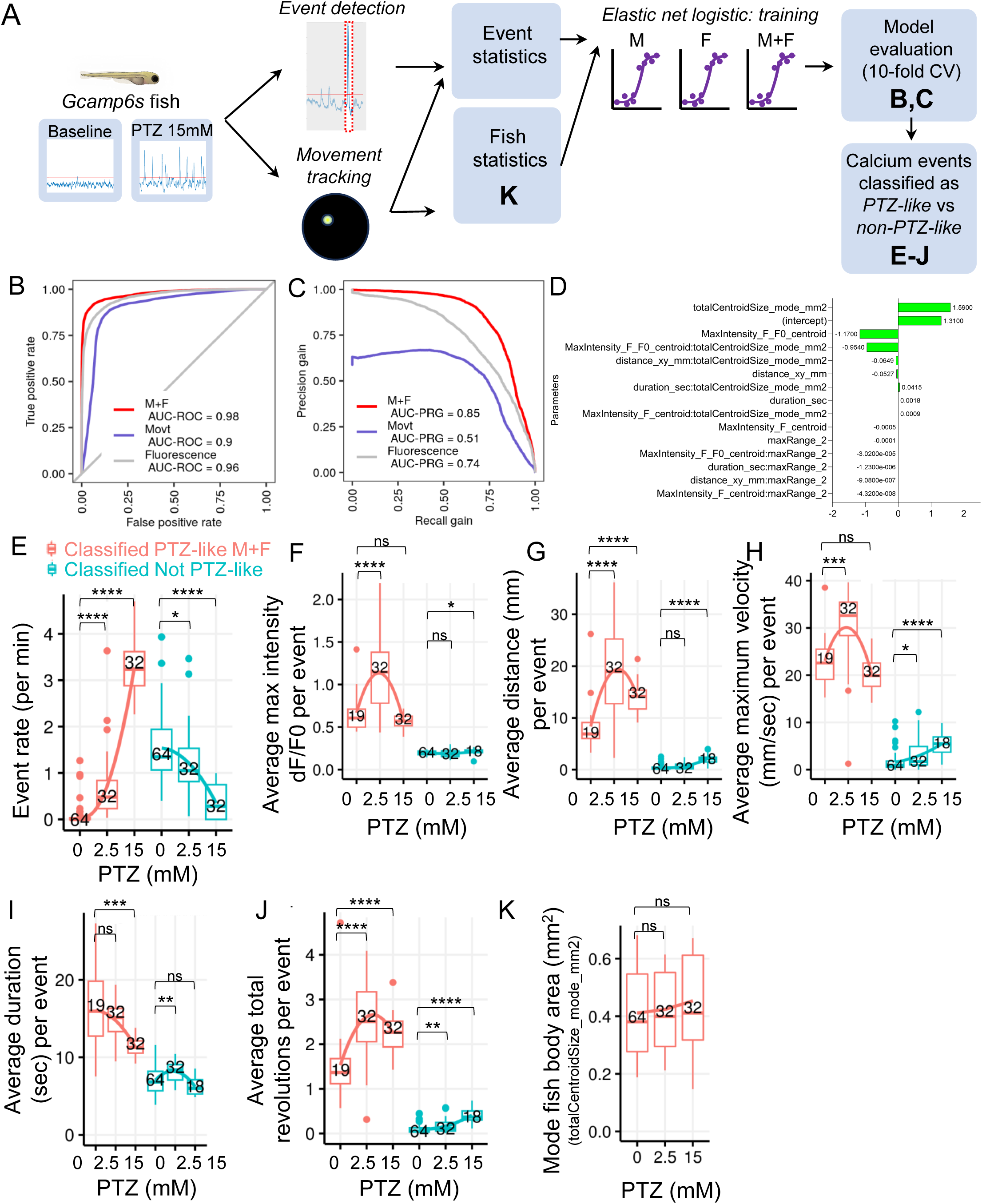
Elastic net logistic regression classifies events as seizure-like with high accuracy. (**A**) Schematic for work-flow to train logistic regression model. Models were trained on either movement data only (M, Movt), fluorescence data only (F, Fluorescence), or combined movement + fluorescence data (M+F). Bold letters indicate figure panels relevant to the indicated step. (**B-C**) Performance of 3 classifiers (M+F, Movt, Fluorescence). (**D**) Parameter tuning for the PTZ M+F model shows features related to fish brightness (“totalCentroidSize_mode_mm2”), peak magnitude of deltaF/F0 during events (“MaxIntensity_F_F0_centroid”), and the interaction between these two features (“MaxIntensity_F_F0_centroid: totalCentroidSize_mode_mm2”) are weighted highly. **(E-J)** Characteristics of events classified as “seizure-like” (*red*) versus “not seizure-like” or physiological (*blue*) by M+F classifier for different doses of PTZ (0, 2.5, 15mM) in the serial dose escalation paradigm. Data are groupwise boxplots of per-fish averages of the indicated event-level parameter, a portion of which was previously used for training. (**K**) Fish body area, a surrogate measure of fish brightness (referred in the model as “totalCentroidSize_mode_mm2”) is not significantly different following PTZ dose escalation. Data are groupwise boxplots of per-fish measures in Tukey format. The number of larvae per group is indicated as numerals; variations in group number between PTZ levels and figures is related to whether fish had any detectable events of the indicated type. Statistical testing is FDR-adjusted pair-wise Wilcoxon rank sum test. *, adj P<0.05. **, adj P<0.01. ***, adj P<0.001. ****, adj P<0.0001.

We asked whether predictors derived from movement features (“Movt”), fluorescence-based features (“Fluorescence”), or a combination of both (“M+F”) would yield the model with the highest accuracy (see **Methods** for model formulae). In lieu of manually labeling each event, we designated all events from baseline recordings as physiological (or “non-PTZ-like”) calcium discharges, while all events from PTZ 15mM treatment were designated “seizure-like”. Data were divided by 70:30 train:test split and models trained to distinguish seizure-like events vs non-seizure like events using 10-fold cross-validation. Model performance was evaluated as the average across all folds, with a hold-out data set available to assess over-fitting.

We find that all three models (Movt; Fluorescence; M+F) perform well (**Fig 3B-C**, **Table 1**) though the M+F model is superior (AUC-ROC 0.98; F1 0.912) versus the Fluorescence model (AUC-ROC, 0.96; F1, 0.87), which in turn is superior to the Movt model (AUC-ROC, 0.9; F1, 0.842). In addition, there are notable differences between the models with respect to precision-recall gain (**Fig 3C**), with the movement-only model demonstrating clearly inferior precision gain (related to the ‘positive predictive value’) across all levels of recall gain (AUC-PRG, 0.51), whereas the fluorescence-only model (AUC-PRG, 0.74) tracks the performance of the M+F model more closely (AUC-PRG 0.85). Nevertheless, the superiority of the M+F model over the alternative models is evidence that fluorescence and movement-based measures are complementary or non-redundant in assigning the seizure-like classification. Lastly, the results of parameter tuning from the M+F model demonstrate that the features weighted most highly by the elastic net regression procedure include a per-fish measure of fish brightness, a per-event measure of event intensity, as well as the interaction between those two features (**Fig 2D**), suggesting that the model is accounting for differences in the seizure-related magnitude of dF/F0 fluctuations that relate to individual differences in the brightness of each fish. Additional performance metrics of the models are in **Table 1**.

**Table 1.** Performance of logistic classifiers trained on combined movement and fluorescence measures (**M+F),** movement only (**Movt**), or fluorescence only (**Fluorescence**). SENS, sensitivity. SPEC, specificity. MCC, Matthew’s Correlation Coefficient. PREC, precision (positive predictive value). NPV, negative predictive value. FPR, false positive rate. AUC-ROC, area-under-curve receiver-operator curve. PR, precision-recall. PRG, precision-recall gain.

|  | **M+F** |  |  | **Movt** |  |  | **Fluorescence** |  |
| --- | --- | --- | --- | --- | --- | --- | --- | --- |
|  | *Value* | *CI* |  | *Value* | *CI* |  | *Value* | *CI* |
| **SENS** | 0.915 | 0.91-0.92 |  | 0.887 | 0.88-0.89 |  | 0.868 | 0.86-0.87 |
| **SPEC** | 0.939 | 0.94-0.94 |  | 0.85 | 0.84-0.85 |  | 0.916 | 0.91-0.92 |
| **MCC** | 0.854 |  |  | 0.727 |  |  | 0.784 |  |
| **Informedness** | 0.855 |  |  | 0.736 |  |  | 0.783 |  |
| **PREC** | 0.908 | 0.9-0.91 |  | 0.802 | 0.79-0.81 |  | 0.871 | 0.86-0.88 |
| **NPV** | 0.944 | 0.94-0.95 |  | 0.916 | 0.91-0.92 |  | 0.913 | 0.91-0.92 |
| **FPR** | 0.061 |  |  | 0.15 |  |  | 0.084 |  |
| **F1** | 0.912 |  |  | 0.842 |  |  | 0.87 |  |
| **AUC-ROC** | 0.98 | 0.98-0.98 |  | 0.9 | 0.9-0.9 |  | 0.96 | 0.96-0.96 |
| **AUC-PR** | 0.97 |  |  | 0.77 |  |  | 0.95 |  |
| **AUC-PRG** | 0.85 |  |  | 0.51 |  |  | 0.74 |  |

To better understand the properties of seizure-like events selected by the classifiers, we computed per-fish averages for each of the two classes of events (seizure vs non-seizure) identified by the M+F model during PTZ dose escalation (0, 2.5mM, and 15mM) and reviewed differences in the group-wise distributions across a variety of measures (**Fig 3E-I**).

First, we observe that the average rate of events (**Fig 3A**) classified as seizure-like is minimal during the baseline state (0.0766 +/- 0.23 per min) – consistent with the goal for the PTZ M+F classifier to select against physiological or spurious events. Second, we observe a dose-response in the rate of seizure-like events relative to PTZ concentration, consistent with our observations from representative data (**Fig 1C**) and from the literature^17^. The detection of seizure-like events in the low-dose PTZ group suggests that some calcium events share qualities with the events contained in the high-dose PTZ training set, but occur at lower rates. Third, the rate of events classified as non-seizure are highest during the baseline state and lowest during high-dose PTZ, also consistent with the behaviors which the classifier is meant to exclude. Fourth, the group-wise average for a key fluorescence measure (**Fig 3F**) used in the model (max intensity dF/F0 per event) comports with our representative data, with highest F/F0 values seen during low-dose PTZ (2.5mM) and still elevated but lower dF/F0 values observed during high-dose PTZ (15mM). This reduction in F/F0 with higher PTZ likely reflects an artifact of fish movement or the moving average deltaF/F0 method, but this evidently does not compromise the accuracy of the M+F classifier or our ability to discern differences in the rate of seizure-like events that differ by PTZ dose. Meanwhile, the group-wise average F/F0 values for events classified as physiological never exceeds 0.3 dF/F0. Fifth, the group-wise distributions of multiple movement-based measures including average distance per event (**Fig 3G**), maximum velocity (**Fig 3H**), total revolutions per event (**Fig 3J**) are elevated following PTZ treatment in events classified as seizure-like, and are uniformly low in events classified as non-seizure. Sixth, the duration of seizure-like events is elevated following PTZ (>10 seconds) but remains brief among non-seizure events (< 10 seconds) (**Fig 3I)**. Seventh, we see no effect of PTZ dose-escalation on the group-wise mean of a measure of fish body area estimated from fluorescence intensity data (**Fig 3K**), suggesting that this measure is reasonably stable across conditions despite escalating seizure-like activity and suitable as a fish-level covariate in the model. Without directly performing a model sensitivity analysis, these results demonstrate the range of parameters typical for events classified as seizure-like activity under the PTZ M+F model.

Lastly, we examined the performance of our classifiers on calcium events related to caffeine treatment (**Supplementary Fig 4**) – a compound reported to increase locomotor activity in larval zebrafish but without known proconvulsant properties at the concentrations tested (0.1 mM)^28^. First, fish exposed to caffeine travel greater distance versus controls (**Supplementary Fig 4B**; p=0.008, Mann-Whitney), consistent with its reported effect. Second, we observe that events classified as seizure-like following caffeine exposure are rare using the M+F classifier (**Supplementary Fig 4Bii**) and not significantly different versus vehicle treated animals (p=1.0, Mann-Whitney test) as predicted. However, events classified as seizure-like by the movement-only classifier are elevated substantially above background though not different from vehicle, suggesting false positive classifications in both groups due to elevated locomotor activity.

Taken together, the M+F model out-performs alternative models based on fluorescence or movement alone, and yields interpretable fish-level estimates of seizure-like activity due to the proconvulsant PTZ.

### 3.3. Combined movement and fluorescence profiling detects differential effects of ASMs in larval zebrafish

We next asked whether combined movement and fluorescence profiling could detect responses to antiseizure medication treatment.

We first tested the response to valproic acid (VPA, 5mM) – an ASM with known anti-seizure activity in larval zebrafish PTZ assay by behavioral locomotor^5^, tectal electrographic^5^ and calcium fluorescence measures^17^ as well as activity in the rodent PTZ assay by behavioral^5^ and electrographic^5^ read-outs. During a serial PTZ dose escalation (**Fig 4A**), we examined the effect of prolonged (20 hours) vs. acute (2 hours) VPA pre-treatment on features of calcium events and the rate of seizure-like activity, with and without the classifiers.

**Figure 4.**
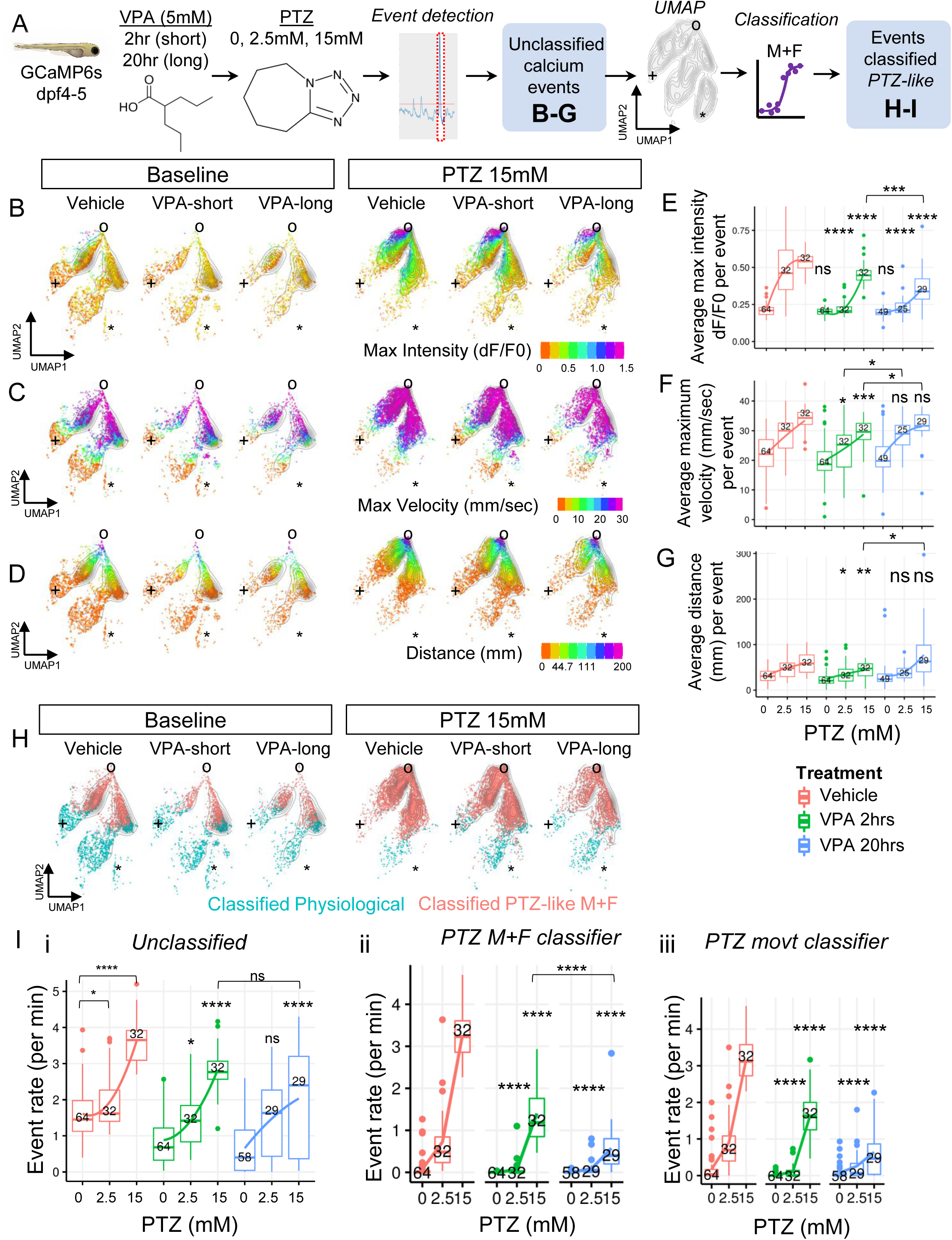
Valproic acid (VPA) treatment differentially alters features of calcium events and reduces overall seizure frequency. (**A**) Overview of experiment to assess effects of acute (2hr) versus prolonged (20hr) pre-treatment with VPA 5mM on PTZ-induced seizure-like activity in PTZ dose escalation. (**B-G**)UMAP of calcium events and their group-level quantification stratified by fluorescence intensity (B, E), maximum velocity (C, F), and distance traveled (D,G). (**H**) UMAP of calcium events classified as seizure-like using movement and fluorescence features (PTZ M+F classifier) (**I**) Rate of PTZ-related calcium events without classification (i), or following classification with PTZ M+F classifier (ii) or PTZ Movt classifier (iii). UMAP data are colored according to indicated color scales. Fiducial markers (+,o,*) relate to the common event embedding for all data reported in this study. Boxplots are Tukey format. The number of larvae per group is indicated as numerals; variations in group number between PTZ levels and figures is related to whether fish had any detectable events of the indicated type. Statistical testing is FDR-adjusted pair-wise Wilcoxon rank sum test. *, adj P<0.05. **, adj P<0.01. ***, adj P<0.001. ****, adj P<0.0001. n.s., not significant. Results of statistical tests are reported relative to control at corresponding PTZ dose, unless otherwise indicated with bars.

First, we observed that both acute and prolonged VPA treatments suppress the PTZ-related increase in the average per-event maximum intensity of normalized calcium fluorescence (dF/F0) (**Fig 4B,E**). The response was most dramatic at low-dose PTZ, where the maximum calcium fluorescence intensity after either acute or prolonged VPA remained similar to baseline controls. At high-dose PTZ, the effect was more mild with prolonged VPA showing greater effects than acute VPA, but both significantly reduced relative to control. Second, movement-related features including average per-event maximum velocity and distance travelled were mildly reduced at low and high dose PTZ with acute VPA, but not with prolonged VPA (**Fig 4C-D**, 4F-G). This may suggest that movement-related features of calcium events are more susceptible (relative to max intensity dF/F0) to sedative effects of acute ASM treatment, since these differences are not present after prolonged VPA, even though the effect of prolonged VPA is stronger than acute VPA with respect to reductions in dF/F0. These findings underscore the increased sensitivity that fluorescence profiling adds to the evaluation of drug responses in larval zebrafish, even before classification is applied.

We next assessed the rate of detected calcium events with VPA treatment before and after applying classification. First, without classification or filtering, both acute and prolonged VPA treatments significantly suppressed the PTZ-related increase in the rate of calcium events at high-dose PTZ (**Fig 4I.i**), though the magnitude of the effect appears similar and there is no evidence for differences between acute vs. prolonged treatment. Second, at low-dose PTZ, only acute VPA suppresses what is observed to be a very mild PTZ-related increase in events (**Fig 4I.i)**, whereas prolonged VPA is associated with a broad range of event rates that are not significantly different. These observations suggest that an approach to classify calcium events may enhance the ability to assess the anti-seizure response of ASMs.

Applying the PTZ M+F classifier, we observe that both acute and prolonged VPA suppressed PTZ-related seizure-like events at both low- and high-dose PTZ (**Fig 4I.ii**). In addition, a robust dose-response between acute versus prolonged VPA was observable at high-dose PTZ. We also applied the PTZ Movt classifier (Fig I.iii) and saw similar results, suggesting that a small set of locomotor parameters of seizure-like events (maximum velocity and distance) are sufficient to detect the anti-seizure effects of VPA, consistent with the literature^5^.

We next tested the response to tiagabine (TGB, 100uM) – an ASM with known anti-seizure activity in the larval zebrafish PTZ assay by tectal electrographic recordings^5^ and in the rodent PTZ assay^5^ by behavioral read-outs, but with reportedly no anti-seizure activity on the behavioral response of PTZ-treated zebrafish larvae^5^. We reasoned that the response to TGB would be an excellent test for the potential of the combined movement and fluorescence profiling approach, as the dual nature of the platform may yield insights into both locomotor activity and surrogate readouts of neuronal activity, and thus speak directly to the observed discrepancy between behavioral and electrographic effects of TGB in zebrafish. We therefore examined the effects of prolonged TGB (20hrs) on properties of calcium events and the rate of seizure-like activity, with and without the classifiers.

First, similar to VPA, we observe a very strong reduction in the PTZ-related increase in average per-event maximum intensity of normalized calcium fluorescence (dF/F0) at both low- and high-dose PTZ (**Fig 5B,E**). Second, among movement-related features, we see a discrepancy in the effects of TGB on average per-event maximum velocity (**Fig 5C,F**), where TGB dramatically suppressed the effect of PTZ-related increased velocity at low-dose PTZ (p<0.0001) but had no effect at high-dose PTZ. By contrast, TGB strongly suppressed the effect of both low- and high dose PTZ on average per-event distance traveled (**Fig 5D,G**). These results again highlight the complex differences in the effects of anti-seizure drugs on properties of PTZ-related seizure-like activity that are accessible to this approach before event classification is employed.

**Figure 5.**
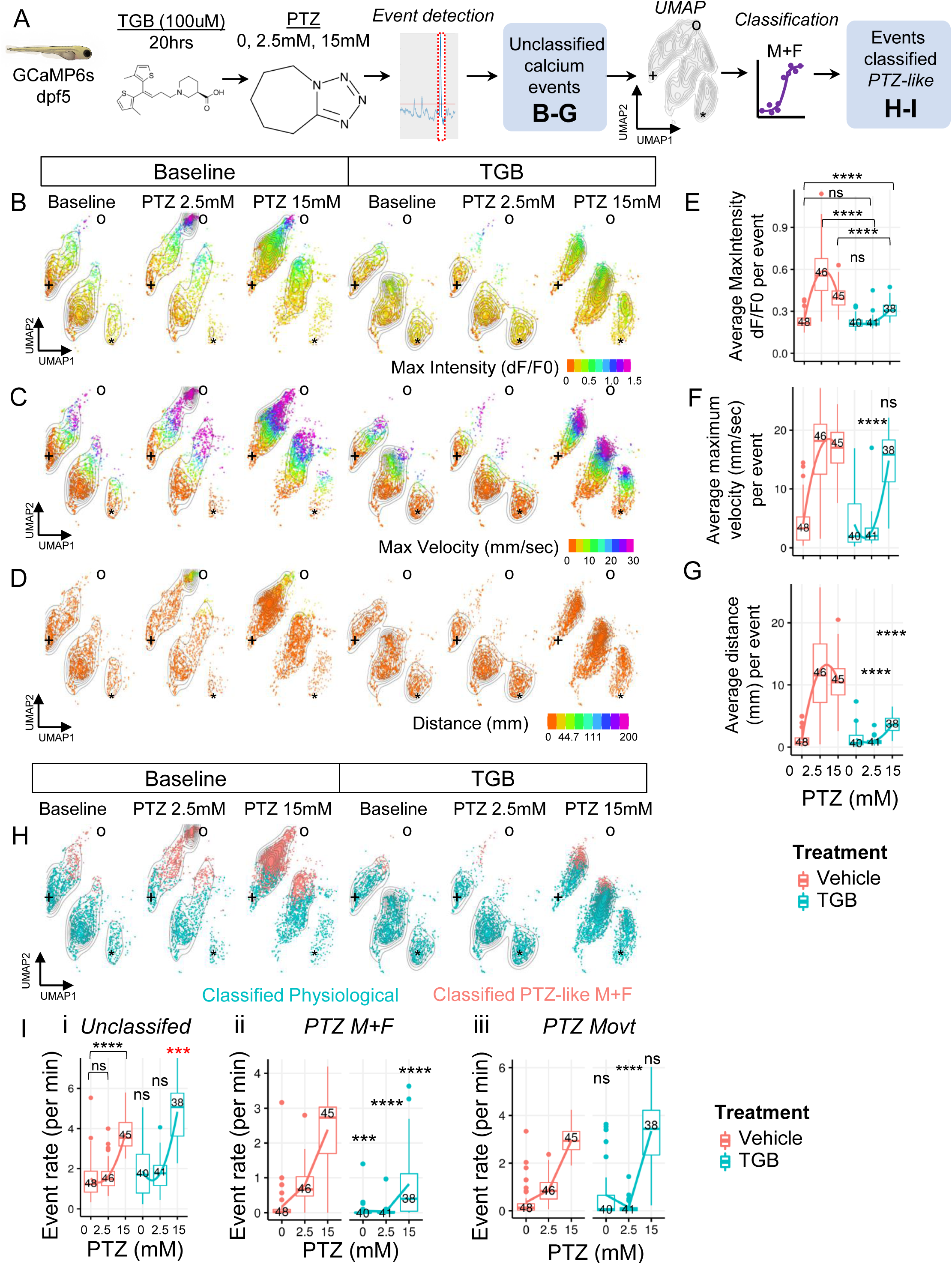
Tiagabine (TGB) suppresses seizure-like activity by differentially altering calcium fluorescence, but not rapid swimming, at high-dose PTZ. (**A**) Overview of experiment to assess effect of prolonged TGB (100uM) pre-treatment on PTZ-induced seizure-like activity in PTZ dose escalation. (**B-G**) UMAP of calcium events and their group-level quantification stratified by fluorescence intensity (B, E), maximum velocity (C, F), and distance traveled (D,G). (**H**) UMAP of calcium events classified as seizure-like (PTZ M+F classifier) (**I**) Rate of PTZ-related calcium events without classification (i), or following classification with PTZ M+F classifier (ii) or PTZ Movt classifier (iii). UMAP data are colored according to indicated color scales. Fiducial markers (+,o,*) relate to the common event embedding for all data reported in this study. Boxplots are Tukey format. The number of larvae per group is indicated as numerals; variations in group number between PTZ levels and figures is related to whether fish had any detectable events of the indicated type. Statistical testing is FDR-adjusted pair-wise Wilcoxon rank sum test. *, adj P<0.05. **, adj P<0.01. ***, adj P<0.001. ****, adj P<0.0001. n.s., not significant. Results of statistical tests are reported relative to control at corresponding PTZ dose, unless otherwise indicated with bars.

We next assessed the effect of TGB on the rate of calcium events before and after applying classification. First, in this cohort of animals, the effect of low-dose PTZ on the rate of unclassified events was negligible, precluding the ability to detect an anti-seizure effect of TGB under these conditions. Second, at high-dose PTZ, TGB was associated with a rate of events that is paradoxically higher than control (**Fig 5I.i**).

Applying the PTZ M+F classifier, TGB dramatically suppressed PTZ-related seizure-like events at both low- and high-dose PTZ (**Fig 5I.ii**), as well as suppressing the low level of events classified as seizure-like at baseline. Interestingly, when applying the PTZ movement classifier, we observe a discrepancy in the effect of TGB, with a dramatic reduction in seizure-like activity at low-dose PTZ (**Fig 5I.iii**) but no effect at high-dose PTZ. Both classifiers therefore identify a significant elevation in the rate of seizure events in response to low-dose PTZ in untreated animals, and a significant suppression of seizures in response to prolonged TGB. However, between the two classifiers, the discrepancy at high PTZ appears to be driven by model term “maximum velocity”, which is one of only two terms included in the “PTZ Movt” classifier. The term is not included in the “PTZ M+F” classifier, highlighting how different classifiers may be tuned to different aspects of the PTZ response.

One interpretation of these results is that although the rate of calcium events with high maximum velocity (consistent with movement criteria for seizure-like activity) remains elevated after high-dose but not low-dose PTZ, the effect of TGB is to reduce the severity of these events through reductions in other measures included in the model. Indeed, we find that fish brightness (**Supplementary Fig 5B**), event duration (**Supplementary Fig 5C**), average velocity (**Supplementary Fig 5D**), and maximum fluorescence per-event (**Supplementary Fig 5E**) are all significantly reduced in unclassified calcium events following TGB treatment, confirming reduced severity of PTZ-related events in these fish. As anticipated, these findings appear to account for the reported discrepancy between the anti-seizure activity of TGB against PTZ in zebrafish tectal LFP but lack of activity in zebrafish locomotor assays^5^, underscoring how combined movement and fluorescence profiling extends the utility of behavioral assays.

In summary, the above method offers new advantages for larval zebrafish PTZ assays with an expanded profile of ASM responsiveness compared to methods based solely on locomotor activity, and enhanced throughput compared to fluorescence microscopy or tectal electrophysiology.

### 3.3. Machine-learning enables low replicate whole organism screening for anti-seizure compounds or targets

We next asked whether our approach would be suitable for low replicate screening for novel anti-seizure interventions. In a primary screen for anti-seizure compounds or genetic targets, there is a need to balance testing as many interventions as possible on as few animals as necessary to identify true hits while limiting false positive results^29^. To explore the range of replicate samples that may be employed in our assay to maximize statistical power while limiting the false positive rate (FPR), we performed a series of simulation experiments using nonparametric bootstrap resampling^30^ and the robust strictly standardized mean difference (RSSMD, or SSMD*^22^) statistic (**Fig 6A-B**, and **Methods**) to assess the optimal RSSMD threshold for defining an antiseizure response in treated animals based on differences in the seizure event rate as a function of bootstrap sample size. The RSSMD is a measure of effect size and variability commonly employed in high-throughput screening, similar to z score^22^.

**Figure 6.**
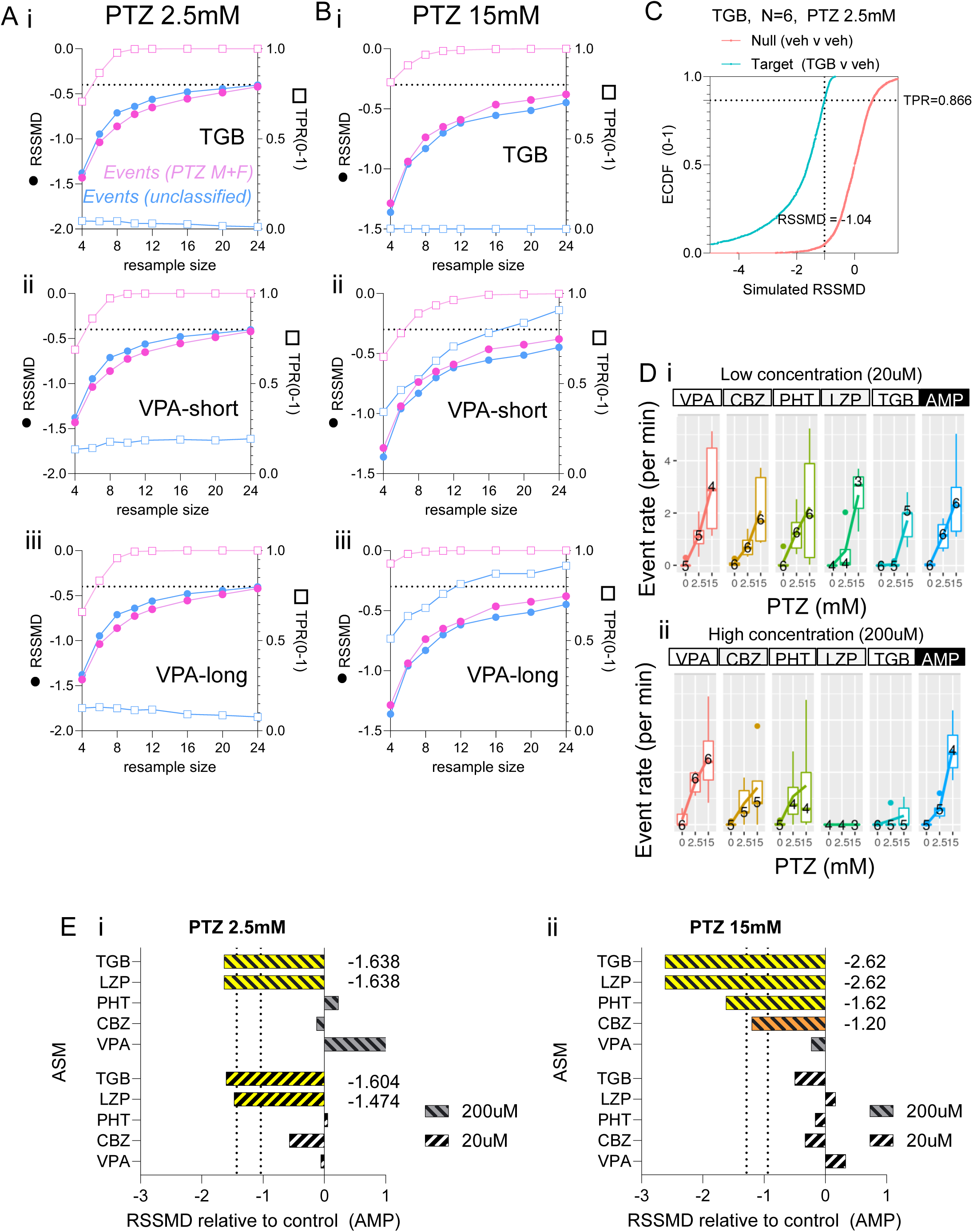
Machine-learning enables low replicate whole organism screening for anti-seizure compounds or targets. **(A-B)** Bootstrap resampling simulations (3000 iterations with replacement) from low-dose PTZ (A) and high-dose PTZ (B) for each ASM dataset, TGB (i), acute VPA (ii), and prolonged VPA (iii). For each bootstrap sample size N (x-axis), closed circles (left y-axis) are the RSSMD threshold required to limit the false positive rate (FPR) to 5% and open boxes (right y-axis) are the associated true positive rate (TPR). Dashed reference lines indicate 80% TPR. (**C**) A representative example of the bootstrap procedure used to generate each data point in (A-B). (**D**) Effect of known anti-seizure drugs (VPA, CBZ, PHT, LZP, TGB) versus negative control (AMP, ampicillin) at low dose (20uM, *i*) or high dose (200uM, *ii*) on PTZ-related seizure-like activity. Data are box-plots of rate of events classified as PTZ-like using the PTZ M+F classifier from multiple fish. N per group is indicated as a numeral on the graph. (**E**) RSSMD values for each ASM relative to on-plate negative control (AMP) at low-dose PTZ (i) and high-dose PTZ (ii). Dashed lines indicate RSSMD thresholds for each PTZ-level and sample size: For PTZ 2.5mM, RSSMD threshold: -1.4309 (N=4), -1.038 (N=6). For PTZ 15mM, RSSMD threshold: -1.286 (N=4), -0.938 (N=6). Colors indicate hits beyond RSSMD threshold: yellow (N=4), orange (N=6). VPA, valproic acid. CBZ, carbamazepine. PHT, phenytoin. LZP, lorazepam. TGB, tiagabine. AMP, ampicillin.

Briefly, for each antiseizure drug, bootstrap resampling distributions of RSSMD values were computed for a range of bootstrap sample sizes (4 - 24; 3000 resamples, with replacement) based on random draws from the target (antiseizure drug treated, PTZ+ASM) dataset vs. vehicle treated (no antiseizure treatment, PTZ+VEH) dataset; these distributions are representative of the range of RSSMD values that could be realized in response to anti-seizure drug treatment. A set of null distributions were computed in the same manner, but with RSSMD values computed for separate random draws from the vehicle treated dataset (PTZ+VEH); this distribution represents the range of RSSMD values that could be realized purely by random chance. For simplicity, we merged vehicle treated animals from separate VPA and TGB datasets. For each bootstrap distribution, the RSSMD threshold is chosen as the 5th percentile of the null distribution, corresponding to a 5% FPR; the true positive rate (TPR) is computed based on the percentile of the target distribution corresponding to the RSSMD threshold. A representative example of this procedure for TGB is shown in Figure 6C. We computed the RSSMD screening threshold and TPR for each antiseizure dataset (TGB, acute VPA, prolonged VPA) at low-dose (**Fig 6A**) and high dose PTZ (**Fig 6B**) using the event rate with or without the PTZ M+F classifier.

First, at low-dose PTZ, the use of the PTZ M+F classifier enabled robust screening potential, with bootstrap sample size as low as N=6 routinely achieving greater than 80% TPR across data sets (e.g. TPRN=6= 0.866 (TGB), 0.861 (acute VPA), and 0.833 (prolonged VPA)). Second, we observe that the PTZ M+F classifier is crucially necessary for the assay to succeed on low-dose PTZ, as the use of unclassified event rates failed to achieve a true positive result above 40% under any sample size tested from any antiseizure data set (**Fig 6A.i-iii**), consistent with the weak differences observed in unclassified event rates under these conditions (cf **Fig 4I.i**, **Fig 5I.i**). The proposed RSSMD screening thresholds using classified events at low-dose PTZ are: -1.4309 (N=4), -1.038 (N=6), -0.8609 (N=8).

At high dose PTZ, we observe that the effects of TGB (**Fig 6B.i**) and prolonged VPA (**Fig 6B.iii**) are even more robust, readily achieving greater than 80% TPR even at N=4. Meanwhile, acute VPA does not exceed 80% until N=6-8 (TPRN=6 = 0.78, TPRN=8 = 0.89). Second, the use of unclassified event rates yields mixed results at high-dose PTZ that differ by ASM. For acute or prolonged VPA, unclassified event rates do eventually achieve TPR>80% at N=20 (**Fig 6B.ii**), and N=12 (**Fig 6B.iii)**, respectively), notably worse than with the use of classified event rates. Meanwhile, for TGB, unclassified event rates remain at 0% across the range of bootstrap values tested, consistent with our earlier observations. The proposed RSSMD screening thresholds using classified events at high-dose PTZ are: -1.286 (N=4) -0.938 (N=6), -0.737 (N=8).

Taken together, we conclude that 4-8 replicates may be reasonable for detecting compounds or mutants with anti-seizure properties similar to our positive control ASMs using the combined movement and calcium fluorescence profiling method and classification based on machine-learning.

### 3.4. Prospective validation of ASM screening strategy confirms that few biological replicates are sufficient to detect anti-seizure response

We next asked whether our screening paradigm would be successful at detecting the anti-seizure response of multiple known ASMs (valproic acid (VPA), carbamazepine (CBZ), phenytoin (PHT), lorazepam (LZP), and tiagabine (TGB)) using the proposed screening parameters. We chose to test known antiseizure compounds at 2 concentrations (20 uM, and 200 uM) irrespective of known therapeutic or lethal/toxic dose ranges, as it mirrors the concentrations accessible in the setting of compound screening and recognizes the infeasibility of determining the maximum tolerated dose (MTD) prior to screening. We chose ampicillin as an active negative control – a compound with no known anti-seizure or proconvulsant properties in larval zebrafish at the concentration tested. We performed the assay using a PTZ dose escalation (0, 2.5mM, 15mM) as described above and observed the event rates from the PTZ M+F classifier (**Fig 6D**) as well as the RSSMD (**Fig 6E**) relative to ampicillin for each ASM x concentration x PTZ combination.

First, we observed significant reductions in seizure-like activity in response to 2 of 5 ASMs (TGB, LZP) across 3 of 4 drug concentration X PTZ combinations (ASM 20uM x PTZ 2.5mM; ASM 200uM x PTZ 15mM; and ASM 200uM x PTZ 15mM; **Fig 6Ei-ii**). Second, the combination of high-dose PTZ (15mM) and high drug concentration (200uM) showed significant reductions in 4 of 5 ASMS (TGB, LZP, PHT, and CBZ; **Fig 6Eii**). Third, despite establishing the anti-seizure efficacy of VPA at higher doses (5mM) in earlier experiments, no anti-seizure effect was observed for VPA at the lower concentrations employed in this experiment, consistent with the known potency of VPA in literature^5,31^.

These findings demonstrate differential pharmacoresponsiveness of PTZ-related seizure activity that differs by PTZ dose. In addition, for screens of novel compounds at a single concentration, our findings suggest that the high-dose PTZ paradigm may provide a greater yield for detecting anti-seizure responses specifically when high test concentrations (200uM) are employed.

## 4. Discussion

The aim of this study was to establish a data-driven platform for higher throughput evaluation of seizure-like activity in larval zebrafish, making use of combined movement and calcium fluorescence in order to assess the effects of anti-seizure interventions. Such an approach would enhance the current offerings for the evaluation of seizure-like activity in zebrafish by integrating two of the most well studied readouts in a single assay, without sacrificing throughput or requiring custom hardware.

We report success in achieving this objective through the use of a commercially available fluorescent plate reader (FDSS7000EX) and seizure-like event classification using a type of supervised machine learning known as elastic net logistic regression^25^.

Our approach to profile calcium fluorescence from unrestrained fish using a fluorescent plate reader is novel, but other authors have reported previous attempts to combine recordings of movement and neural activity using lower through-put approaches. For example, Turrini et al. reported use of a head-fixed / tail-free configurations^17^ to record simultaneous brain calcium fluorescence and high-speed tail movement during PTZ exposure. Another report employed technically sophisticated custom hardware to dynamically reposition and refocus a microscope objective to track movement and acquire neuroluminescence (not calcium fluorescence) from a single unrestrained fish^32^. The main reason that our approach succeeds despite allowing movements of the fish (which likely contribute in part to fluctuations in measured calcium fluorescence) and despite imaging from the ventral surface of the fish (which reduces signal strength compared to dorsal imaging) is likely due to our narrow use of the platform to study seizure-like activity. The robust signals associated with synchronized neuronal discharges during seizure-like events are acquired relatively easily, and rather than “solving” issues related to variation in fish brightness or movement, we show how machine learning provides a tractable data-driven approach to classification in the face of these complexities.

### 4.1. Features of calcium events mirror established classification scheme for seizure-like activity in larval zebrafish

Use of the GABA_A_R antagonist PTZ as a pharmacological model of absence epilepsy in rodents^33^ and as a general proconvulsant in zebrafish^4,5^ is extensive and well-established. Historically, pharmacological models of PTZ-induced seizures have reported specific behavioral, electrographic, and pharmacoresponsive profiles^33^. Behaviorally, the rodent PTZ model shows dose-specific findings with motionless staring occurring at low-dose PTZ, clonic or myoclonic seizures arising at intermediate-dose PTZ and tonic seizures arising at higher doses^33^. In larval zebrafish, PTZ-induced seizures have been observed to follow a sequence of severity from early hyperactivity (Stage 1), bouts of rapid circular swimming (Stage 2), followed by tonic contraction, clonic convulsions, and motionless loss of posture (Stage 3). However, the time-averaged prevalence of these grades at different PTZ-dose levels has not been systematically reported. Electrographically, rodent models demonstrate generalized spike and wave discharges on EEG, very similar to human EEG findings^33^, while tectal recordings of larval zebrafish show periodic interictal discharges and higher amplitude ictal discharges^4,5,11,13^.

Here we show that combined movement and fluorescence profiling identifies event features similar to those reported in the literature for locomotor assays of PTZ-induced seizures. At low-dose PTZ (2.5mM), most calcium events are classified as “physiological”, but there is an increase in seizure-like events over baseline, comprised of rapid circular swimming, elevated distance travelled and high dF/F0. This is likely comparable to Stage 1/2. At high-dose PTZ (15mM), seizure-like calcium events heavily predominate over the physiological type, which may be similar to Stage 2/3.

### 4.2. Machine learning

We show that the PTZ M+ F classifier trained with elastic net logistic regression has the highest performance metrics, compared to fluorescence only or movement only classifiers, and demonstrate through bootstrap simulations that its use in our screening approach enables relatively low numbers of biological replicates.

The use of machine learning to classify the behavior of larval zebrafish under different experimental conditions has been previously reported^13,34,35^, but the present approach employed several innovations including the use of combined fluorescence and movement data, the use of elastic net logistic regression, as well as event-level (as opposed to subject-level) classification.

Our use of elastic net regression with interactions was necessitated by the variation introduced by calcium fluorescence, stemming from intrinsic differences in fish brightness that persisted despite pre-screening fish based on GCaMP6s brightness (which is a common procedure in lower throughput methods) and normalization of calcium fluorescence using the common deltaF/F0 approach. Given excellent results with elastic net regression, we did not systematically investigate alternative ML approaches and it is possible that other methods could produce similar or superior results. However, our preliminary analyses did suggest that “elastic net” was superior to unpenalized methods, which demonstrated poor generalizability across different cohorts of fish (data not shown), likely due to lack of regularization. In addition, the use of elastic net enables straightforward interpretation of model tuning to understand the relative weighting of different features after model training, compared to methods based on deep-learning architectures. Lastly, classification at the event-level was an early design choice in order to quantify the seizure susceptibility of individual fish, which has the advantage of yielding continuous data at the fish-level which is more interpretable and statistically tractable than fish-level categorical classifications.

### 4.3. Differences in effect of VPA vs TGB

We report differences in the responsiveness of high-dose PTZ-related calcium events to VPA versus TGB treatment, with VPA reducing seizure-like activity using either the PTZ M+F or PTZ Movt classifier, whereas TGB selectively shows reductions only with the PTZ M+F classifier. Both classifiers show reductions to VPA or TGB at low-dose PTZ.

Although VPA is an ASM with known anti-seizure activity in the larval zebrafish PTZ assay by behavioral locomotor^5^, tectal electrographic^5^ and calcium fluorescence measures^17^, TGB instead has only been shown to have activity by tectal electrographic assay and with reportedly no anti-seizure activity on the behavioral response of PTZ-treated zebrafish larvae^5^. As noted previously, we reasoned that the response to TGB would test the potential of the combined movement and fluorescence profiling approach to yield insights into both locomotor activity and surrogate readouts of neuronal activity, and address the observed discrepancy between behavioral and electrographic effects of TGB in zebrafish.

Indeed, it appears that our platform with the two classifiers is suitable for detecting these distinct assay-dependent responses. These findings highlight the potential of combined movement and fluorescence profiling to augment both traditional locomotor-based and neurophysiological assays.

Our choice of features for the PTZ Movt classifier (distance, max velocity) was meant to incorporate the features most frequently reported as useful for detecting seizure-like activity on locomotor assays^4,5,11–13^. By comparison with the PTZ M+F classifier, our goal was to highlight conditions under which methods that are limited to locomotor behavior may perform similarly or differently to combined movement and fluorescence profiling. However, the results from one instantiation of a “movement based” classifier should not suggest that TGB treatment does not alter the movement of PTZ-treated larvae. Indeed, although we reported that the maximum velocity of unclassified calcium events at high-dose PTZ was unchanged by TGB, we report several other event features that are reduced by TGB including distance travelled and mean velocity.

Furthermore, it is worth noting that the performance of the PTZ Movement classifier is respectable, consistent with the strong evidence basis for using locomotor assays to assess the response of PTZ in larval zebrafish^4,5^ . Although we did not formally test the following assumption, it is possible that the use of the PTZ Movt classifier or a similar approach based on the rate of seizure-like movement bouts could improve the statistical performance of locomotor assays performed on other platforms (e.g. Noldus Ethovision), which traditionally report “distance travelled” or “integrated activity units” in consecutive time bins.

### 4.4. Screening strategy validation

Our prospective validation of the ASM screening strategy reveals differences in the pharmacoresponsiveness of PTZ-related seizure activity that may differ by dose.

Pharmacologically, based on work established in rodent models^33^, models of absence epilepsy including PTZ-induced seizures demonstrate responsiveness to certain classes of antiseizure medication (ethosuximide, valproic acid, benzodiazepines) and exacerbation to other classes (for example sodium channel blockers such as phenytoin, carbamazepine; barbiturates; as well as GABA_A_ receptor agonists (gaboxadol) and GABAB receptor agonists (such as baclofen, and GHB).

The zebrafish PTZ model overall has demonstrated similar patterns of responsiveness, including: 1) responsive to benzodiazepines (CZP^4^, DZP^4,5^); 2) response to VPA^4,5^; and 3) exacerbation by GABAB receptor agonists (Baclofen^4^). However, there are also notable differences in the zebrafish PTZ model versus rodent PTZ. First, there is conflicting evidence regarding sodium channel blockers. PHT has been reported to have no effect on locomotor or EEG seizures^5^, but it has also been reported to reduce EEG seizures^4^. CBZ has been reported to have no effect on locomotor^5^ or EEG seizures^4,5^, but improvements on both locomotor and EEG have also been reported^36^. To review, the effect of sodium channel inhibition should be to exacerbate seizures, at least based on rodent PTZ data and human patients with generalized epilepsy^33^. Second, the T-type calcium channel blocker ethosuximide has been reported to reduce locomotor and EEG seizures^5^, but one report showed no effect on EEG seizures^4^. Lastly, the response to GABA_A_R agonists is also anomalous in zebrafish. Phenobarbital has been reported to reduce EEG seizures^4^, whereas no effect of primidone (a drug similar to phenobarbital) was observed on locomotor and EEG seizures^5^. Again, both compounds exacerbate generalized seizures in rodent and humans^33^. In the context of our results, we demonstrate the efficacy of the benzodiazepine, LZP, and GAT1 inhibitor, TGB, at both low- and high-dose PTZ, which is consistent with the literature.

In addition, we report that our ASM screening strategy is suitable for detecting the response of 4 out of 5 known ASMs, including efficacy of sodium channel inhibitors CBZ and PHT selectively in the setting of high-dose PTZ exposure. Although CBZ and PHT are not historically recognized to be active in PTZ models^33^, and reportedly exacerbates discharges in rodent models^33^, the zebrafish literature has conflicting evidence with most reports showing no effect on PTZ-related locomotor activity (actineg units)^5^ or tectal EEG abnormalities^4,5^, but with at least one report showing efficacy of CBZ against PTZ-related activity, based on locomotor assay of movement counts and midbrain EEG activity in larval zebrafish^36^.

One interpretation may be that low-dose PTZ in our assay may be the more faithful screening concentration, as it confirmed the anti-seizure effect of two types of ASMs known to be active against PTZ, and did not identify ASMs that have not previously been confirmed. However, this is unlikely to be the case, given the majority of results in the literature using the equivalent of our high-dose PTZ (15 mM^4^ or 20mM^5^). To the best of our knowledge, although rodent models of PTZ demonstrate dose-dependent behavioral manifestations, there are no previous reports demonstrating dose-dependent differences in pharmacoresponsiveness. Another interpretation is that our method may capture more of the substrates quantified by the approaches employed by authors reporting an anti-seizure effect of CBZ^36^. Settling this matter should be pursued in future investigations.

Taken together, these observations highlight how our M+F approach appears to succeed despite still unexplained discrepancies in the measured efficacy of ASMs to PTZ that differ by modality, compound, and perhaps other factors in larval zebrafish.

Overall, the results of our studies show that it is feasible to perform combined movement and fluorescence profiling with the help of machine learning for the purpose of assaying seizure-like activity and the response to antiseizure interventions in larval zebrafish using a relatively small number of biological replicates. This should allow high throughput screening using the robotic and fluid handling capabilities of modern plate readers such as the FDSS7000EX that we employed. In addition, we anticipate the use of combined movement and fluorescence profiling to support reverse genetic screens such as with the recently reported MICDrop technique^37^ to identify targets whose of loss-of-function cause seizure-like activity or suppress the effects of proconvulsants, the latter of which may serve as substrates for the development of novel anti-seizure small molecules. Our approach is equally applicable to proconvulsant drugs with different pathways of neurotransmission, which have been relatively less studied in larval zebrafish^38,39^, setting the stage for more comprehensive characterization of larval zebrafish models of chemical seizures. Lastly, our approach may also hold promise for the evaluation of zebrafish models of genetic epilepsy, which we report in a separate investigation.

## Data availability statement

All of the data generated in the present study and MATLAB/R code are available upon request.

## Acknowledgements

The authors would like to thank Lee Barrett for technical assistance with the FDSS7000EX, and members of the Poduri lab including Cristina Baker and Christopher LaCoursiere for helpful discussions.

## Author contributions

Conceived and designed the experiments: CM. Performed the experiments: CM

Data analysis: CM

Wrote the manuscript: CM

Revised and approved the final version of the manuscript: AP, CM

---

**Supplementary Figure 1.**
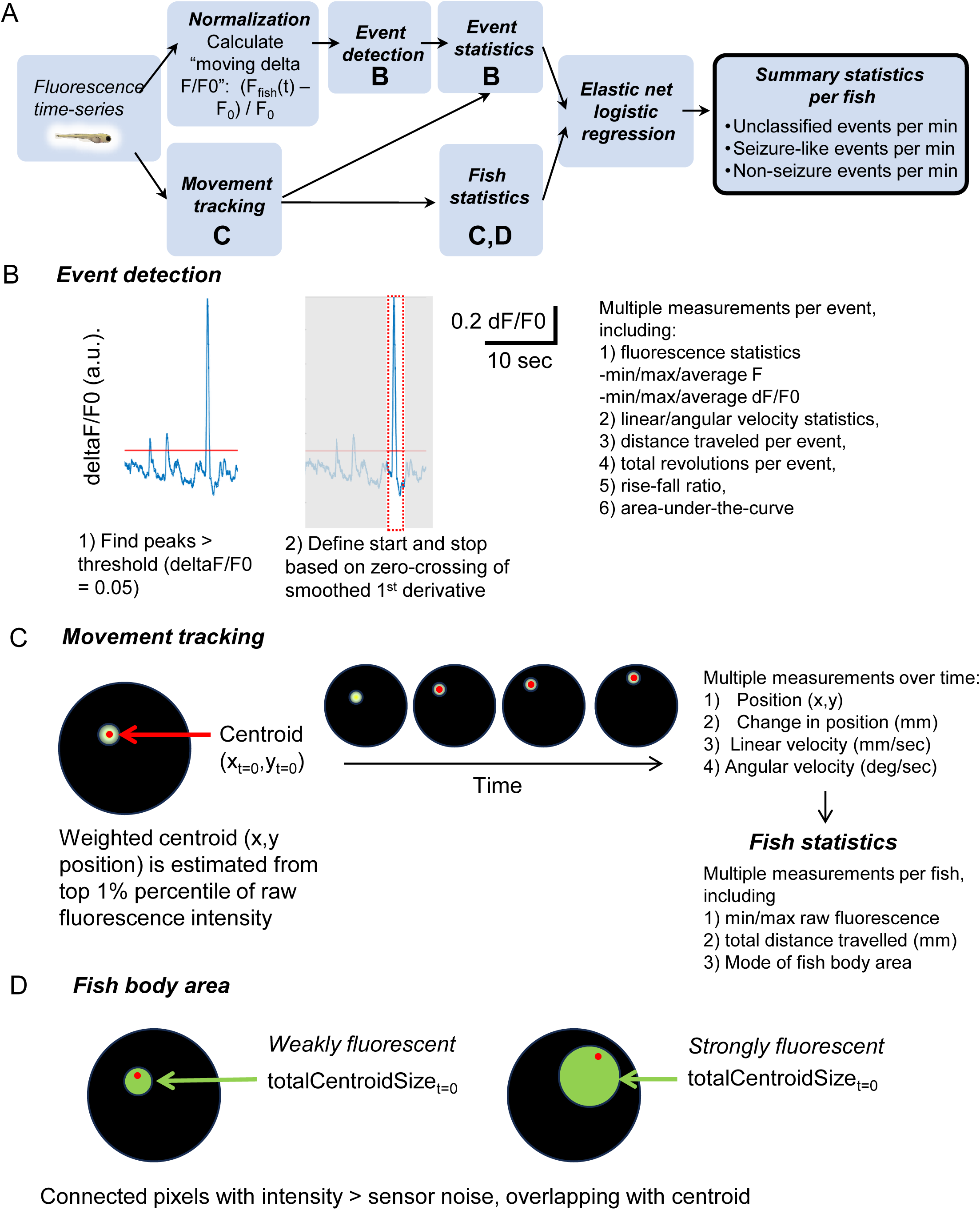
Visual description of method used for combining movement and fluorescence profiling. (A) Overview (B) Event detection (C) Movement tracking (D) Depiction of “fish body area” (total centroid size).

**Supplementary Figure 2.**
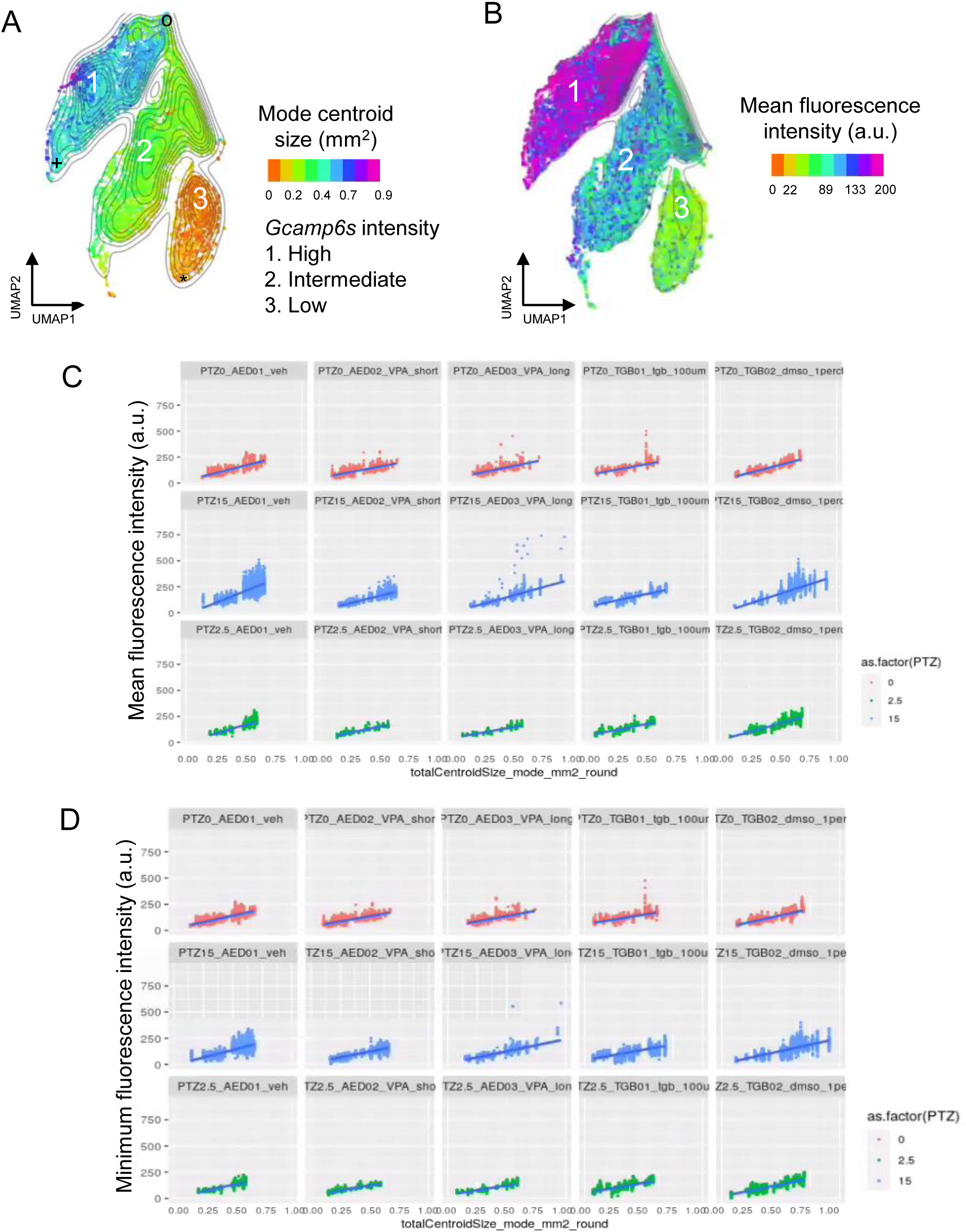
(**A-B**) UMAP demonstrates that centroid size (A) and mean fluorescence intensity (B) are distributed very similarly. Data are calcium events pooled from all fish. (**C-D**) Linear relationship between cross-sectional area of GCaMP6s fluorescence (“totalCentroidSize_mode_mm2”) and average fluorescence intensity (“meanIntensity_F_centroid”) or minimum fluorescence intensity (“meanIntensity_F_centroid”) across conditions and animals. Data values from individual fish

**Supplementary Figure 3.**
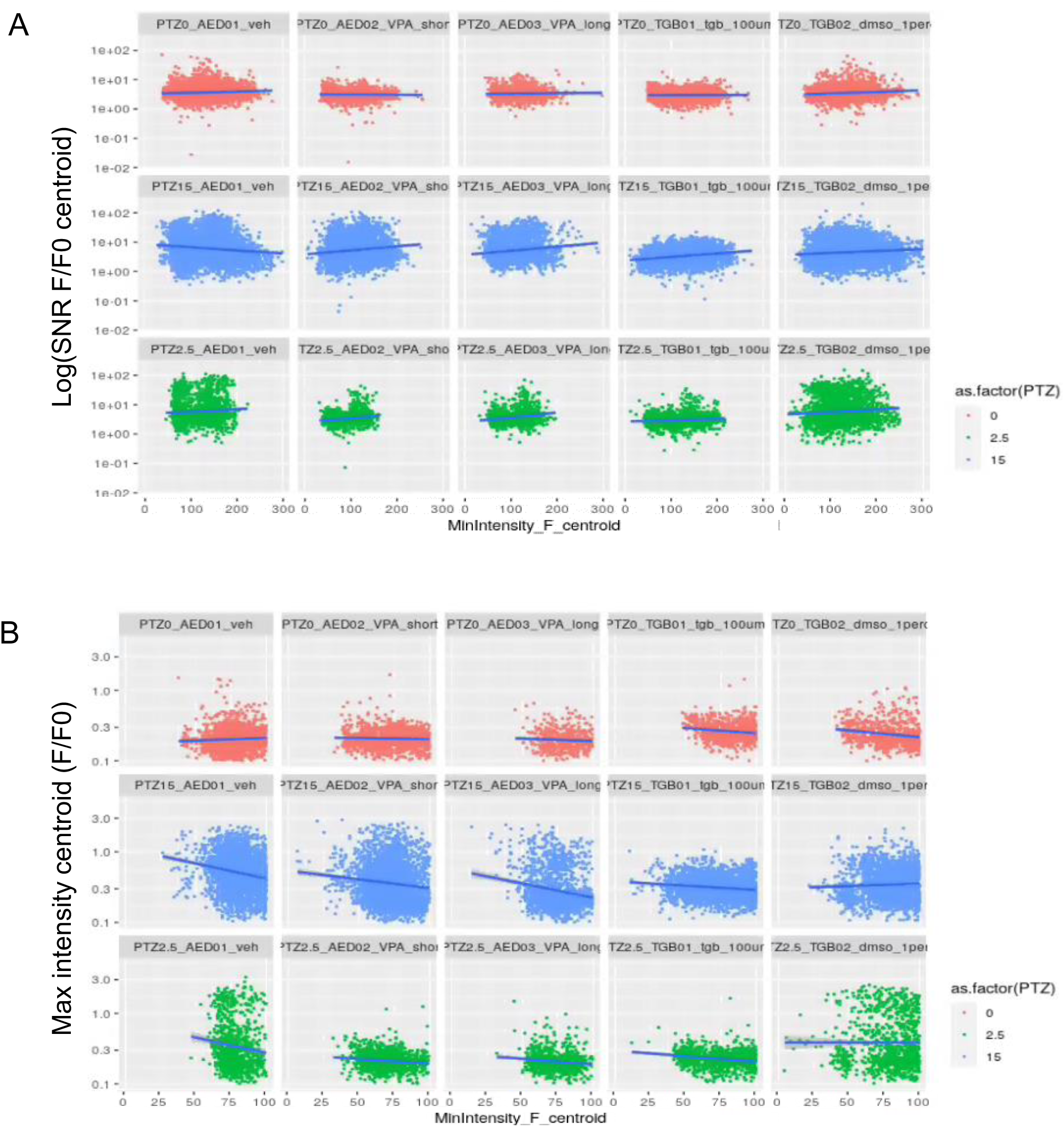
Relationship between minimum fish brightness and SNR_dF/F0 or max intensity dF/F0. Data values from individual fish.

**Supplementary Figure 4.**
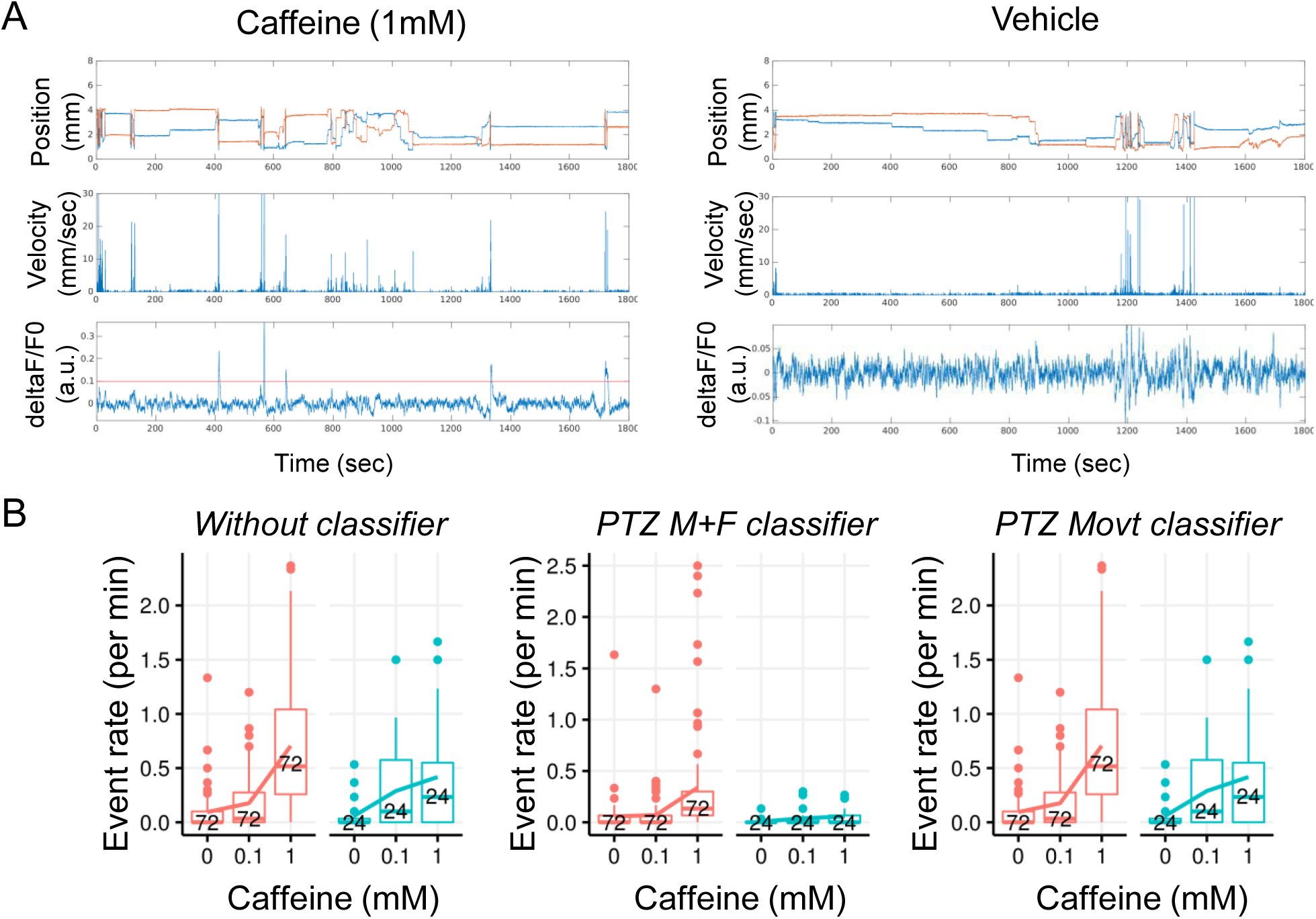
The PTZ M+F classifier excludes non-specific caffeine-induced locomoter activity better than Movt classifier. **A**) Representative FDSS data following administration of caffeine (1mM; left) vs vehicle (H20; right). (**B**) Event rates increase with administration of caffeine (red) vs vehicle (blue) based on unclassified events (i) or with events classified as PTZ-like using the PTZ Movt classifer (iii) but only modestly increase when the PTZ M+F classifier is applied (ii)

**Supplementary Figure 5.**
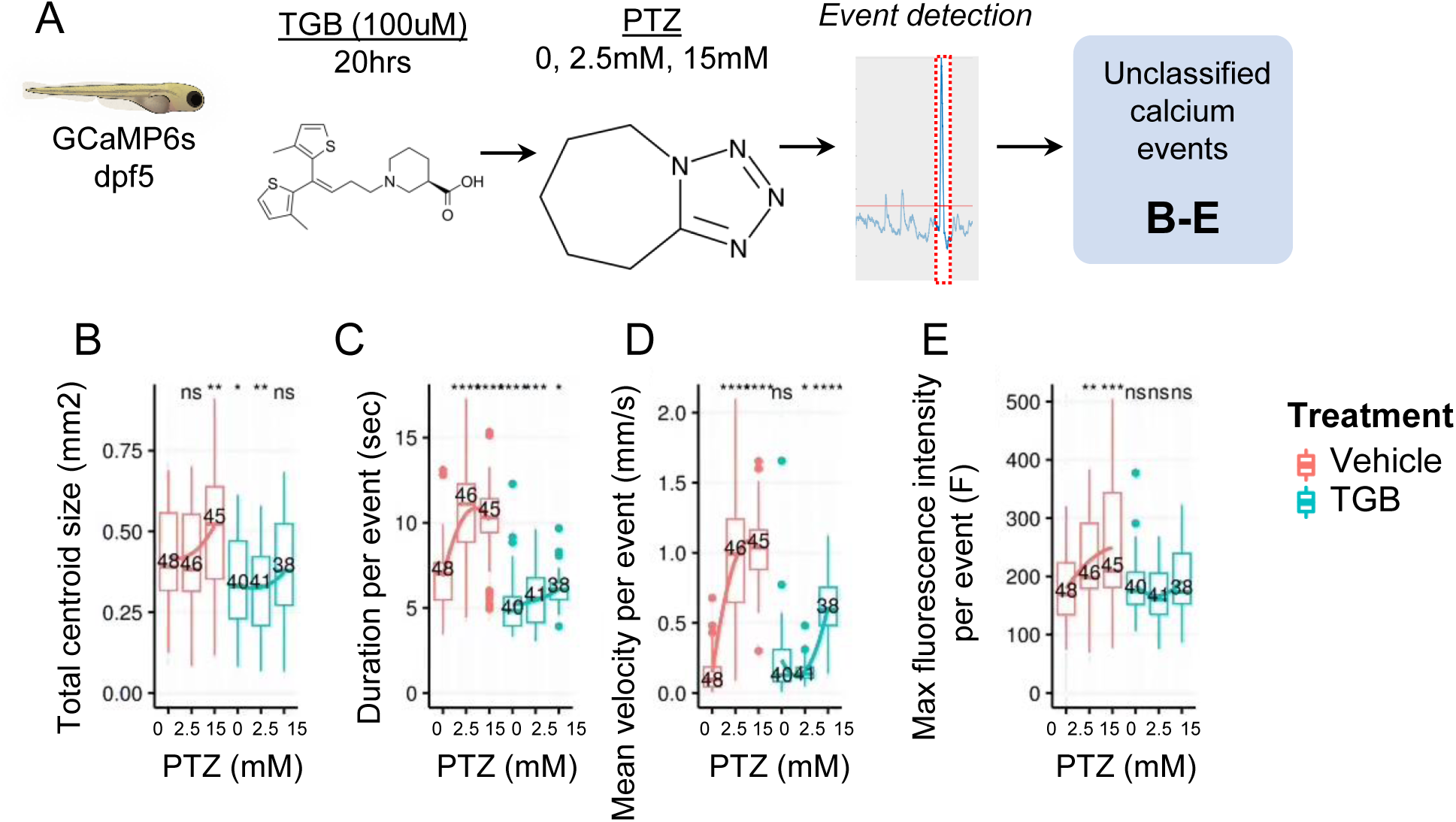
Multiple properties of calcium events are altered by TGB treatment. (**A**) Overview. (**B-E**) Total centroid size (B), duration per event (C), mean velocity per event (D), and maximum unnormalized fluorescence intensity per event (E)

**Supplemental Video 1. Video of 96 unrestrained GCaMP6s larval zebrafish under baseline conditions.** Pseudo-colored fluorescence time-series from 30-minute recording (displayed at ∼2.8X real-time), MP4 compression. File name: FDSS_96wp_01_veh_200923004_raw.mp4

**Supplemental Video 2. Video of 96 unrestrained GCaMP6s larval zebrafish exposed to proconvulsant (PTZ 15mM).** Pseudo-colored fluorescence time-series from 30-minute recording (displayed at ∼2.8X real-time), MP4 compression. File name: FDSS_96wp_03_PTZ15mM_200923006_raw.mp4

